# First virological and pathological study of Göttingen Minipigs with Dippity Pig Syndrome (DPS)

**DOI:** 10.1101/2023.01.26.525667

**Authors:** Hina Jhelum, Nanna Grand, Kirsten Rosenmay Jacobsen, Sabrina Halecker, Michelle Salerno, Robert Prate, Luise Krüger, Yannik Kristiansen, Ludwig Krabben, Lars Möller, Michael Laue, Benedikt Kaufer, Kari Kaaber, Joachim Denner

**Affiliations:** Institute of Virology, Free University, Berlin, Germany; Scantox A/S, DK-4623 Lille Skensved, Denmark; Ellegaard Göttingen Minipigs A/S, Dalmose, Denmark; Marshall BioResources, North Rose, New York, USA; Robert Koch Institute, Berlin, Germany; Robert Koch Institute, Centre for Biological Threats and Special Pathogens ZBS 4: Advanced Light and Electron Microscopy, Berlin, Germany

## Abstract

Dippity Pig Syndrome (DPS) is a well-known but rare complex of clinical signs affecting minipigs, which has not been thoroughly investigated yet. Clinically affected animals show acute appearance of red, exudating lesions across the spine. The lesions are painful, evidenced by arching of the back (dipping), and the onset of clinical symptoms is generally sudden. In order to understand the pathogenesis, histological and virological investigations were performed in affected and unaffected Göttingen Minipigs (GöMPs). The following DNA viruses were screened for using PCR-based methods: Porcine cytomegalovirus (PCMV), which is a porcine roseolovirus (PCMV/PRV), porcine lymphotropic herpesviruses (PLHV-1, PLHV-2, PLHV-3), porcine circoviruses (PCV1, PCV2, PCV3, PCV4), porcine parvovirus 1 (PPV1), and Torque Teno sus virus (TTSuV1, TTSuV2). Screening was also performed for integrated porcine endogenous retroviruses (PERV-A, PERV-B, PERV-C) and recombinant PERV-A/C and their expression as well as for the RNA viruses hepatitis E virus (HEV) and SARS-CoV-2. Eight clinically affected and one unaffected GöMPs were analyzed. Additional unaffected minipigs had been analyzed in the past. The analyzed GöMPs contained PERV-A and PERV-B integrated in the genome, which are present in all pigs and PERV-C, which is present in most, but not all pigs. In one affected GöMPs recombinant PERV-A/C was detected in blood. In this animal a very high expression of PERV mRNA was observed. PCMV/PRV was found in three affected animals, PCV1 was found in three animals with DPS and in the healthy minipig, and PCV3 was detected in two animals with DPS and in the unaffected minipig. Most importantly, in one animal only PLHV-3 was detected. It was found in the affected and unaffected skin, and in other organs. Unfortunately, PLHV-3 could not be studied in all other affected minipigs. None of the other viruses were detected and using electron microscopy, no virus particles were found in the affected skin. This data identified some virus infections in GöMPs with DPS and assign a special role to PLHV-3. Since PCMV/PRV, PCV1, PCV3 and PLHV-3 were also found in unaffected animals, a multifactorial cause of DPS is suggested. However, elimination of the viruses from GöMPs may prevent DPS.

## Introduction

Dippity Pig Syndrome (DPS) is a common clinical term for symptoms that may be diagnosed as acute dermatitis or erythema multiforme [1–10]. The affected animals show acute red, exudating lesions that develop characteristically across the spine. These lesions are painful, evidenced by arching of the back (dipping), and sensitivity to touch. Clinical signs range from the pigs being visibly uncomfortable (e.g, by pacing, not resting), to showing obvious signs of pain, with arching of the back and vocalizing. The onset of clinical signs is often sudden, developing within hours or even minutes. The lesions on the back appear wet with exudation, showing fluid running down the sides in addition to erythema. By the second day, the lesions are typically covered with crust and are in the healing phase. The syndrome is self-limiting (but should always be pain treated), and the lesions will heal in a relatively short period of time (a few days to 2 weeks) with no scarring. In Göttingen minipigs, the syndrome is recognized in all age groups, whereas in potbelly pigs it has been described as more commonly affecting pigs less than one year old [1–6].

In human medicine, erythema multiforme is characterized as a polymorphous erythematous rash confined to the skin [11]. It is considered to be a hyperergic mucocutaneous immune-mediated reaction to infections. In humans, herpes simplex virus (HSV), a human alphaherpesvirus, also called human herpesvirus (HHV), is usually the causative agent in patients with erythema multiforme minus, with HSV-1 (HHV-1) more common than HSV-2 (HHV-2). These viruses are also the cause in the majority of adults suffering from erythema multiforme majus that is characterized by typical target lesions on the extremities. In rare cases erythema multiforme in adults is caused by another herpesvirus, varizella zoster virus (VZV, human herpesvirus 3, HHV-3, or human alphaherpesvirus 3) [12]. In children, adolescents and young adults, a large percentage of erythema multiforme major cases is caused by *Mycoplasma pneumoniae;* here, the lesions are usually atypical and predominantly occur on the body [13]. Other authors think that human erythema multiforme is an acute, immune-mediated condition characterized by distinctive lesions of the skin [14].

Recently, commercial breeding pigs suffering from a syndrome similar to erythema multiforme were analyzed [15, 16]. The animals showed red skin areas, and the disease was acute, self-limiting and often associated with anorexia, fever and respiratory problems. Blood and skin samples were investigated. When screened for different porcine viruses using sensitive methods developed for screening of donor pigs and recipients in xenotransplantation, PLHV-1 was found in affected skin areas of five animals, PLHV-2 was found in one animal and PLHV-3 in four animals. Neither PCMV/PRV, nor PCV1-4 were found in the affected skin. In the blood of these animals PLHV-1 was present only in two animals, indicating that replication is mainly ongoing in the affected skin. PLHV-2, PCV2 and PCV3 were found in the blood of one animal. Noteworthy, in the blood of one animal four different viruses (PLHV-1, PLHV-2, PCV2 and PCV3) were found simultaneously. All animals were positive for PERV-C, but recombinant PERV-A/C was not found [16].

The cause of DPS in minipigs is not known. It is thought that stress may be an important contributing factor [17]. A correlation between DPS and exposure of the animal to the sun has also been considered [9]. DPS can occur as a single, one-time event, or an individual pig can suffer multiple rashes over time. It occurs most often in young pigs – between 4 months and 4 years, and is seem more rarely in older pigs, even though they may have been affected as younger piglets [9]. Dipping or temporary loss of use of hind legs were observed, but DPS usually does not affect front legs. The diseased animals recover within 2-4 days without medical intervention, however, affected minipigs should always be treated for pain [10].

Some believe that DPS and erythema multiforme are one and the same, whereas others suggest that DPS is only similar to erythema multiforme that has long been recognized in commercial pigs [15, 18–20]. It is unclear whether pigs drop their hind end due to the severe pain inflicted by the condition, or whether a possible causative agent also affects the spinal cord motoric nerves of the hind limbs [10].

Although pig and veterinary associations informed in detail about DPS [7–10], there are only very few scientific publications investigating cases of DPS [1–6, 17] and these publications usually only include a short description of the disease. The literature describing erythema multiforme in commercial breeding pig herds is also rare [15, 16]. Since DPS is found in minipigs bred for biomedical purposes [21, 22], there is a great interest to analyze the cause of this disease in order to prevent it effectively [17]. Among other studies, GöMPs have been used as donors in a preclinical xenotransplantation trial transplanting islet cells into non-human primates [23] and it is planned to use GöMPs for pig islet cell transplantation trials in Germany. GöMPs from the Ellegaard Göttingen Minipigs A/S are well studied concerning prevalence of viruses and other microorganisms. They were found free of 89 microorganisms, with very few animals where PCMV/PRV and HEV were found [23–25]. Here we present the results of the first virological and histological studies on GöMPs suffering from DPS.

## Methods

### Animals and sampling

For the virological screening of pigs suffering from DPS, samples were collected from eight GöMPs and sent to the Robert Koch Institute (RKI), Berlin, or later to the Institute of Virology at the Free University (FU), also in Berlin, Germany. The sample panel included whole heparin-, or EDTA-blood, plasma, serum, several organ samples (liver, spleen, stomach, sternum) and skin punches. Skin punches of individual minipigs were taken from three different regions: Affected, exudative skin (hereinafter referred to as “skin A”), border between red-discolored skin and unaffected skin (“skin B”) and skin not affected (“skin C”) (Figure 1). Blue rectangles in figure 1 indicate the collected sample if the animal was terminated and a large skin sample was taken, whereas the green circle indicates smaller biopsies from animals not terminated.

**Fig 1.**
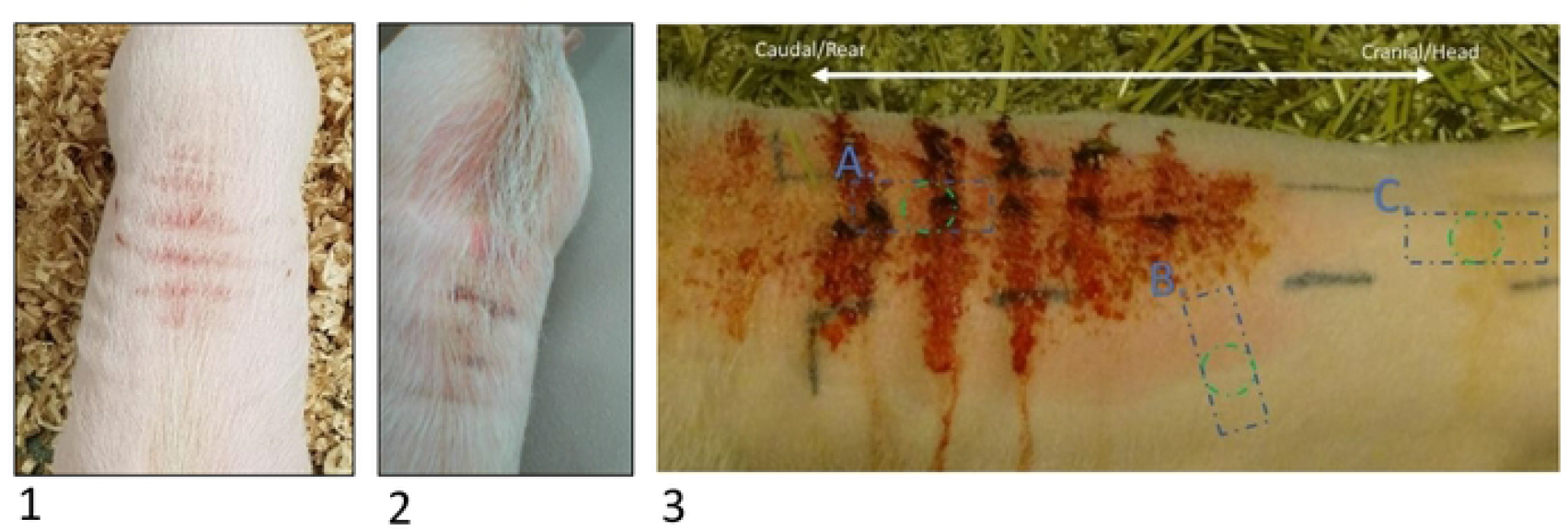
Lower back of GöMPs with acute dippity pig syndrome. Pigs are suffering from red exudative skin alterations exemplary displayed in minipig #343061 (1, 2) and another minipig (3). Samples from different skin areas were included in the virological analyses: Affected skin (A), intermediate skin (B) and unaffected skin (C). Blue squares indicate excisional biopsy, green circles indicate punch biopsy.

Organs (liver, spleen, stomach, sternum) and skin samples from pig #901 and skin samples from pig #239185 were obtained as formalin-fixed, paraffin-embedded (FFPE) tissue sections. Samples from all pigs suffering from DPS were collected either in the acute phase of the DPS or during the healing phase. In addition, blood samples from one pig (#343528) which showed no clinical signs, but was housed close to the affected animal #901, were also studied. No skin samples were obtained from this animal. Additional unaffected GöMPs from Ellegaard Göttingen Minipigs A/S and from the Göttingen University had been screened in the past [24–29].

### Isolation and stimulation of peripheral blood mononuclear cells (PBMCs)

Ethylendiamintetraacetat-treated blood (EDTA-blood) from minipigs #239185 and #237587 was dispensed upon 5 ml Pancoll human (PAN-Biotech GmbH, Aidenbach, Germany) and centrifuged at 900 x g for 21 minutes to isolate PBMCs. They were harvested and washed twice with phosphate buffered saline (PBS) by centrifugation at 350 x g for 10 minutes. 1 x 10^6^ PBMCs per sample were seeded in 12-well plates with 2 ml Roswell Park Memorial Institute medium (RPMI-1640, PAN-Biotech GmbH, Aidenbach, Germany) and stimulated with 2.4 μg/ ml phytohemagglutinin-L (PHA-L) solution (500 x) (Invitrogen, Waltham, MA, USA) for five days. Both, PHA-L stimulated and non-stimulated PBMCs were stored at −20 °C for later processing.

### Nucleic acid extraction from blood, PBMCs and frozen tissues

At the RKI, DNA was extracted from sera, blood or PBMCs from minipigs #342036, #342746 and #343061 using different DNA extraction kits: DNeasy Blood and Tissue kit (Qiagen GmbH, Hilden, Germany), and NucleoSpin Virus (Macherey-Nagel, Düren, Germany). DNA was quantified and the 260 nm/280 nm ratio was determined using a NanoDrop ND-1000 (Thermo Fisher Scientific Inc., Worcester, MA, USA) as described previously [25].

At the Institute of Virology, Free University Berlin, DNA and RNA extraction from minipigs #901, #239185, #237587 and #349753, was performed as follows: Frozen heparin blood was centrifuged at 300 x g for 10 minutes. DNA of the cell pellet was extracted using the Invisorb Spin Universal Kit (Invitek Molecular GmbH, Berlin, Germany) and RNA was extracted using the QIAamp RNA Blood Mini Kit (Qiagen, Hilden, Germany) according to the manufacturer’s instructions. The purified PBMCs and approximately 20 mg of the frozen tissue samples were transferred to innu SPEED Lysis Tubes (Analytik Jena, Jena, Germany) and homogenized twice for 20 seconds with the MP Biomedicals FastPrep-24. DNA and RNA extraction of the homogenized PBMCs as well as the DNA of the homogenized tissues were performed using the innuPREP Virus DNA/ RNA Kit (Analytik Jena, Jena, Germany). RNA extraction from the homogenized tissues was carried out using the RNeasy Lipid Tissue Min Kit (Qiagen, Hilden, Germany).

### Nucleic acid extraction from formalin-fixed paraffin-embedded tissue (FFPE) embedded tissue sections

The QIAamp DNA FFPE Tissue Kit (Qiagen, Hilden, Germany) was used for the DNA extraction of FFPE embedded tissue sections, while RNeasy DSP FFPE Kit was used for RNA extraction of these samples. All protocols were carried out according to the manufacturer’s recommendations.

### Nucleic acid extraction from skin samples

Prior to extraction, the skin was cut into small pieces, transferred to a mix of collagenase (final concentration 125 U/ml, Sigma-Aldrich, St. Louis, MO, USA) and hyaluronidase (final concentration 100 U/ml, Sigma-Aldrich, St. Louis, MO, USA) and incubated for 30 minutes at 37°C and shaking at 500 rpm as recommended by Reimann et al. [30]. After centrifugation for 5 minutes at 1.000 x g, the sample was further processed for nucleic acid extraction. For DNA extraction, DNeasy Blood and Tissue Kit (Qiagen, Hilden, Germany) was used according to the manufacturer’s instructions. To extract the RNA of the pre-incubated samples, the RNeasy Lipid Tissue Mini Kit (Qiagen, Hilden, Germany) was applied and a DNase digestion was carried out by using the RNase-Free DNase Set (Qiagen, Hilden, Germany).

### Real-time reverse transcriptase PCR (real-time RT-PCR) for the detection of HEV

Real-time reverse transcriptase-PCR based on TaqMan technology as described by Jothikumar et al. [31] was carried out to detect hepatitis E virus (HEV). All real-time RT-PCR reactions were prepared in a reaction volume of 16 μl using SensiFAST Probe No-ROX One-Step Kit (Meridian Bioscience, Cincinnati, OH, USA) plus 4 μl template RNA. The experiments were applied as duplex real-time RT-PCRs, thus detecting HEV and an internal control (Influenza A-RNA) per sample [32]. The real-time RT-PCR was performed with the qTOWER^3^ G qPCR cycler (Analytik Jena, Jena, Germany) applying a temperature-time profile that consists of a reverse transcriptase step of 30 minutes at 50°C, followed by an activation step of 15 minutes at 95°C and 45 cycles comprising a step of 10 seconds at 95°C, followed by a step of 20 seconds at 55°C and 15 seconds at 72°C.

### Real-time PCR for the detection of DNA viruses

TaqMan based real-time PCRs were performed to detect PCMV/PRV [33], PLHV-1, PLHV-2, PLHV-3 [34], PCV1, PCV2, PCV3, PCV4 [35, 36] and PPV1 [37] as described previously. PCR detecting the Torque Teno sus viruses 1 and 2 (TTSuV1 and TTSuV2) was established according to [38]. Sequences of the primers and probes are listed in Table 1. All protocols were performed using the SensiFAST Probe No-ROX kit (Meridian Bioscience, Cincinnati, OH, USA) in a reaction volume of 16 μl plus 4 μl of DNA template. Real-time PCRs were carried out as duplex PCRs that simultaneously indicate the gene of interest and porcine glyceraldehyde-3-phosphate-dehydrogenase (pGAPDH) as internal control for each sample. Real-time PCR reactions were carried out with a qTOWER^3^ G qPCR cycler (Analytik Jena, Jena, Germany) and the real-time PCR conditions for the detection of PCMV/PRV, PCV1, PCV4, PLHV-1 and PPV1 were applied as previously described [16]. An adapted PCR-time profile for PCV3 started with an activation step of 5 minutes at 95°C, followed by 40 cycles comprising a denaturation step of 15 seconds at 95°C, an annealing step of 60 seconds at 56°C and an extension step of 30 seconds at 72°C. The temperature-time profiles for TTSuV1 and TTSuV2 were set as follows: 2 minutes at 50°C (activation step), 10 minutes at 95°C (denaturation step), followed by 45 cycles of 15 seconds at 95°C (annealing step) and 60 seconds at 60°C (elongation step).

**Table 1.**
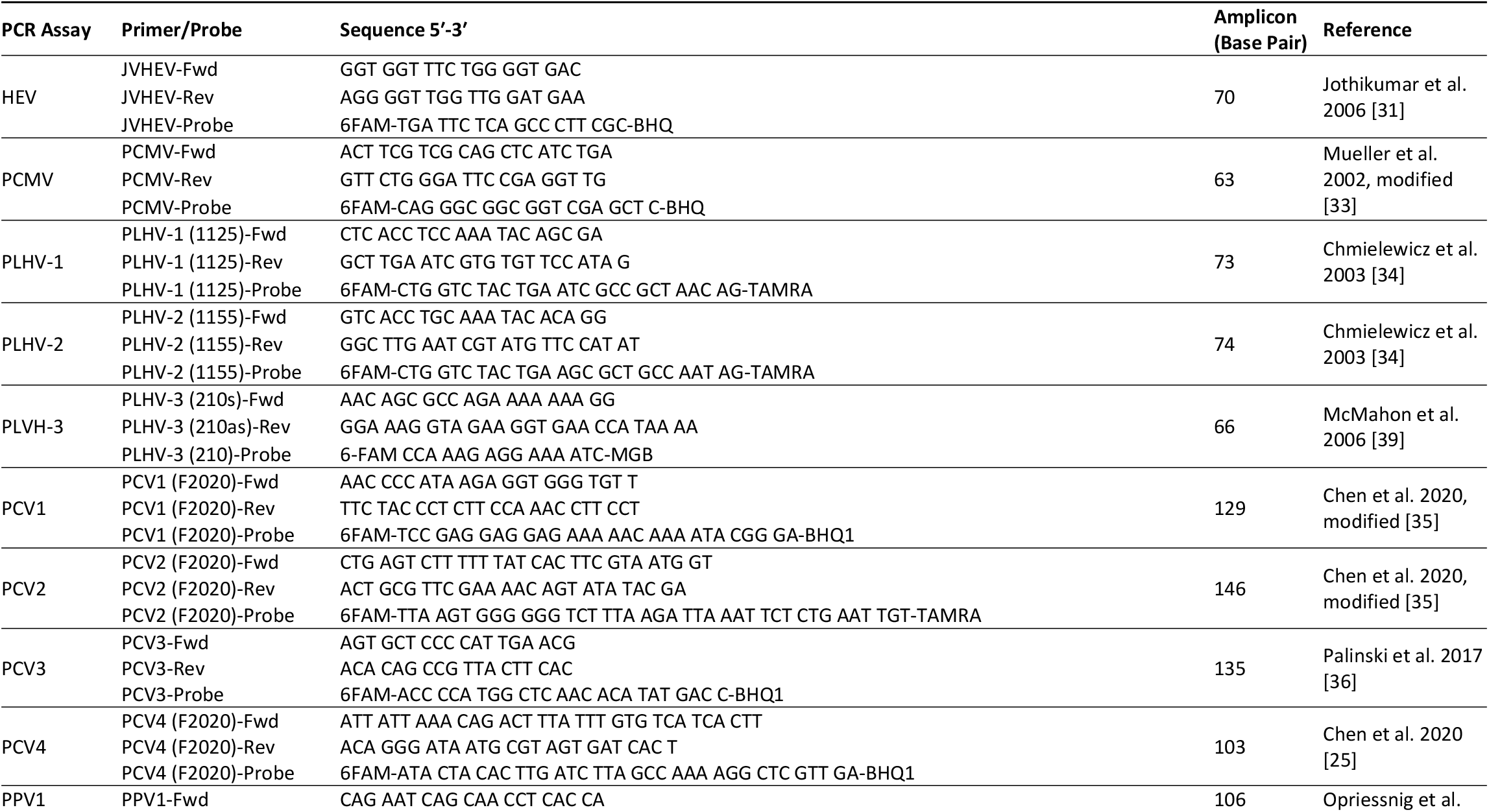
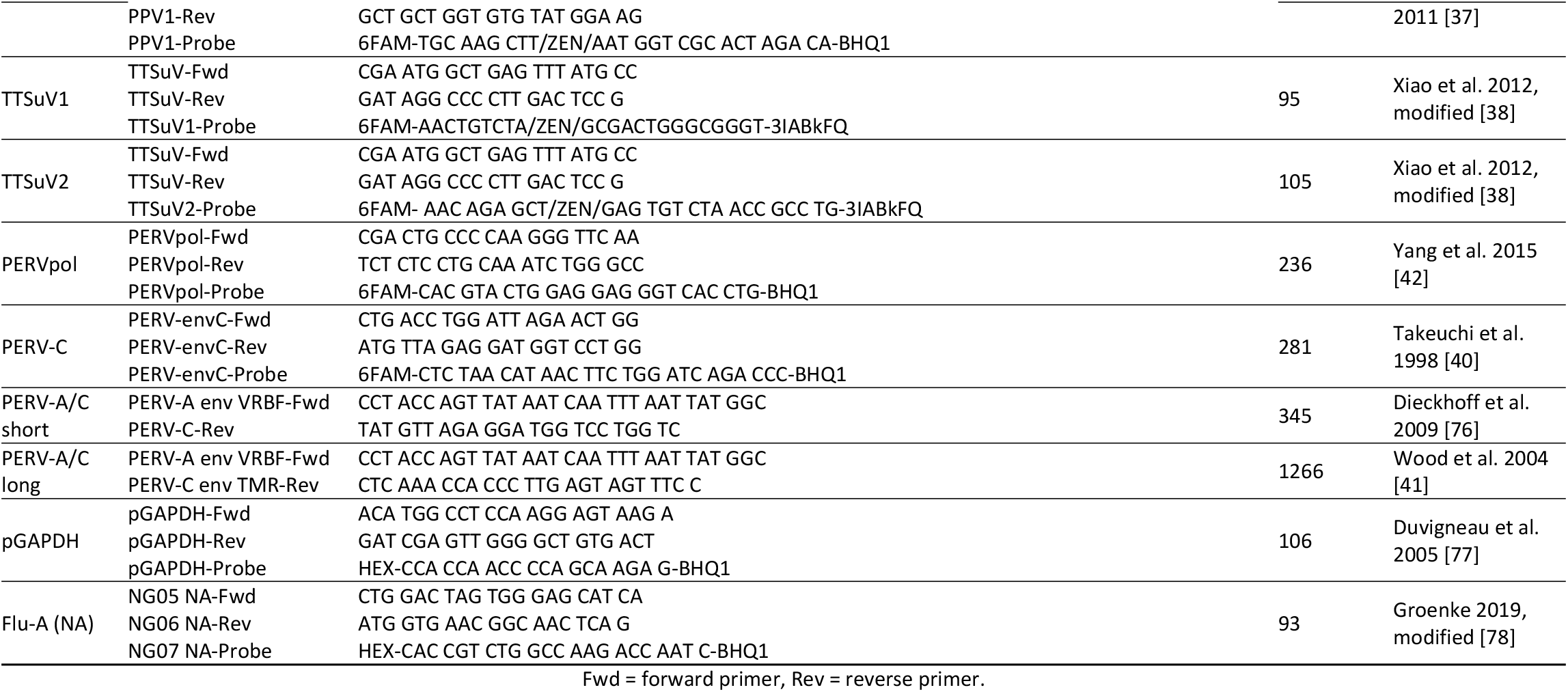
Oligonucleotides for the primers and probes used in this study.

### Real-time RT-PCR for the detection of PLHV-3 mRNA

The real-time RT-PCR was carried out according to McMahon et al. [39]. All reactions were prepared in a volume of 16 μl using SensiFAST Probe No-ROX One-Step Kit (Meridian Bioscience, Cincinnati, OH, USA) plus 4 μl template RNA. Experiments were performed as duplex real-time RT-PCR, thus detecting PLHV-3 and an internal Influenza A-RNA control as described [32]. Real-time RT-PCR was performed with a qTOWER3 G qPCR cycler (Analytik Jena, Jena, Germany) applying a temperature-time profile that consists of a reverse transcriptase step of 30 minutes at 50°C, followed by an activation step of 10 minutes at 90°C and 45 cycles comprising a step of 30 seconds at 90°C, followed by a step of annealing and extension of 30 seconds at 59°C.

### PCR and RT-real-time PCR for the detection of porcine endogenous retroviruses (PERVs)

A PERV-C specific real-time PCR based on TaqMan technology [40] was carried out using the SensiFast Probe No-ROX kit (Meridian Bioscience, Cincinnati, OH, USA). Real-time PCR was set in 16 μl approach plus 4 μl DNA template. The PCR run was performed with a qTOWER^3^ G qPCR cycler (Analytik Jena, Jena, Germany) as follows: Inactivation step at 95°C for 2 minutes followed by 45 cycles comprising a denaturation step at 95°C for 15 seconds and a combined annealing-extension step at 54°C for 30 seconds. The fluorescence signals were measured at the combined annealing-extension step.

A conventional PCR to determine the presence of human-tropic PERV-A/C, was set up using two primer pairs comprising different amplicon lengths (Table1) [41]. The PERV-A/C short primer mix detects an amplicon of 345 base pairs (bp) length, while the PERV-A/C long primer mix detects an amplicon of 1266 bp length (Table 1). Both conventional PCRs were carried out with tAmpliTaq DNA Polymerase (Applied Biosystems, Waltham, MA, USA) and were set up with a Biometra TRIO cycler (Analytik Jena, Jena, Germany). The PCR approach for the short amplicon size was applied with the following temperature-time profile: 95°C for 10 minutes (activation step), followed by 45 cycles of 95°C for 15 seconds (denaturation), 55°C for 30 seconds (annealing) and 72°C for 30 seconds (extension) and a final single cycle of 72°C for 5 minutes. The same temperature-time profile was used for the detection of the long amplicon size, only the time for the extension step was extended by 90 seconds.

A real-time PCR using primers and probe located in the polymerase gene was carried out for the expression of PERV [42]. The PCR mix was set up with the SensiFAST Probe No-ROX One-Step Kit (Meridian Bioscience, Cincinnati, OH, USA) and reaction was carried out with a qTOWER^3^ G qPCR cycler (Analytik Jena, Jena, Germany). The temperature-time conditions were as follows: 50°C at 30 minutes, 95°C for 5 minutes, 45 cycles of 95°C for 15 seconds, 58°C for 30 seconds and 72°C for 30 seconds.

### Western blot analysis to detect antibodies against PCMV/PRV

Western blot analysis was performed as previously described [43, 44]. Briefly, the immunodominant C-terminal fragment R2 of the gB protein of PCMV/PRV was used as antigen [26]. The R2 fragment of the gB of PCMV/PRV was expressed in *E. coli* BL21 cells using the pET16b vector encoding PCMV-R2 as described in detail [44]. Bacteria were induced with 1 mM isopropyl-β-D-1-thiogalactopyranoside (Roth, Karlsruhe, Germany), harvested, and dissolved in 10 mL 8 M urea, 0.5 M NaCl, 15 mM imidazole, 20 mM Tris pH 7.5. After centrifugation the supernatant was applied to a HisTrap HP column connected to an Äkta Prime Plus system (both GE Healthcare, Chicago, Illinois, USA), and eluted after washing using 500 mM imidazole, 6 M urea, 0.5 M NaCl, 20 mM Tris pH 7.5. Purified R2 protein was characterized by sodium dodecylsulfate polyacrylamide gel electrophoresis (SDS-PAGE). The protein was dissolved in sample buffer (375 mM Tris-HCl, 60% glycerol, 12% SDS, 0,6 M DTT, 0.06% bromophenol blue) and denatured for 5 min at 95 °C prior to electrophoresis. SDS PAGE was run in a Mini-Protean Tetra Vertical Electrophoresis Cell (Bio-Rad Laboratories, Incs., Hercules, CA, USA) using a 17% polyacrylamide gel and the PageRuler prestained protein ladder (Thermo Fisher Scientific, Waltham, USA). The protein was transferred for 100 min to a polyvinylidene fluoride membrane (ROTI PVDF, 8989.1, Roth, Karlsruhe, Germany) by electroblotting (100 mA) using the electroblotting device of peqlab Biotechnologie GmbH. After electroblotting the membrane was blocked for 1h at 4 °C in PBS with 5% non-fat dry milk powder (Roth, Karlsruhe, Germany) and 0.05% Tween 20 (Roth, Karlsruhe, Germany) (PBS-T) (blocking buffer). The membrane was cut into strips and incubated over night at 4°C with diluted sera (1:150) in blocking buffer. Afterwards, washing was performed with 0.05% PBS-T three times for 10 minutes each. Polyclonal goat anti-pig immunoglobulin G (IgG) Fc Secondary Antibody HRP (Invitrogen by Thermo Fisher Scientific, Waltham, USA) was diluted 1:15.000 in blocking buffer and strips were incubated for 1 h at room temperature, followed by three washing steps for 10 minutes each. The signal was detected after 5 min incubation with ECL Western Blotting Substrate (Cytiva, Amersham) with a FUSION-SL 3500 WL imaging device (peqlab Biotechnologie GmbH).

### Next generation sequencing (NGS)

To further investigate the possible causative agent of the DPS in minipigs, two skin punches (skin A and skin C) of approximately 1 cm in diameter from pig #237587 were sent to Anicon Labor GmbH (Höltinghausen, Germany) for metagenomic analysis on an Illumina platform including an rRNA depletion in advance. Bioinformatic data analyzes were also provided by Anicon Labor GmbH (Höltinghausen, Germany).

### Electron microscopy

Ultrathin section electron microscopy of skin samples was performed as described by Laue [45].

### Processing of histopathological tissue samples and light microscopic evaluation

Tissue samples for light microscopy were fixed in phosphate buffered neutral 4% formaldehyde. After fixation, samples were trimmed and processed and the specimens were embedded in paraffin and cut at a nominal thickness of approximately 5 μm, stained with haematoxylin and eosin, and examined under a light microscope. Samples for microscopic evaluation were obtained on different days in relation to appearance of clinical signs. Samples from six minipigs #239185, #314, #901, #349753, #237587, and #342036 were taken on the same day as lesions appeared, samples from pig #237629 were taken on the day after lesions appeared and samples from pig #342746 were taken 16 days after lesions appeared.

## Results

### Pathogenesis and occurrence

Eight Göttingen Minipigs with DPS and one animal without signs of disease were analyzed, the skin from affected animals were examined microscopically. Skin samples from the affected animals included samples from skin showing lesions and samples from unaffected, normal skin. In addition, skin from unaffected animals was examined regularly. In previous studies approximately 40 unaffected GöMPs were screened for different viruses without testing skin samples [23–29]. Affected animals included both male and female GöMPs. Most affected animals showed acute parallel lesions with exudate on the dorsal spine, arching of the back (dipping), for some minipigs vocalization and sensitivity to touch was observed. The lesions and other features of the dippity syndrome appeared usually suddenly (e.g., within 2 hours) and lasted several hours up to several days. The example of the GöMPs at the Marshall facility and at their customers shows how seldom the syndrome was observed, which in the beginning was even classified as erythema multiforme (Table 2). However, underreporting cannot be excluded.

**Table 2.** Dippity pig syndrome cases in GöMP from Marshall

| Year | Customer | Cases |
| --- | --- | --- |
| 2018 | 1 | Reports of increasing numbers of Erythema multiforme, since 2015 |
| 2019 | 2 | May: 3 pigs from 3 orders<br>June: 7 cases from 5 different orders |
|  | 3 | 6 cases from 46 (4 control animals, 5 mild cases), 1 month after arrival |
| 2020 | 4 | 4 cases from 48, 5 months after arrival, 3 animals resolved in 2-3 days |
|  | Marshall facility | One case: pregnant sow 2 weeks pre-partum |

### Histology

Histological examination revealed significant changes in the skin of the affected animals (Figure 2). In general, the lesions from the skin samples collected at the same day were characterized by an acute necrotizing inflammation, including epidermal necrosis, epidermal intra-/extracellular edema, epidermal/subepidermal inflammatory cell infiltration (dominated by neutrophilic granulocytes) and vesicle/vesico-pustular formation. Occasionally, superficial crust and subepidermal haemorrhage and congestion of blood vessels were also seen. No bacteria were seen in any of the affected skin samples nor were any intracellular (intranuclear or intracytoplasmic) inclusion bodies reported in the epithelial cell of the epidermis. Normal skin sampled from the affected animals and from other animals did not show any pathological findings at the microscopic examination

**Fig 2.**
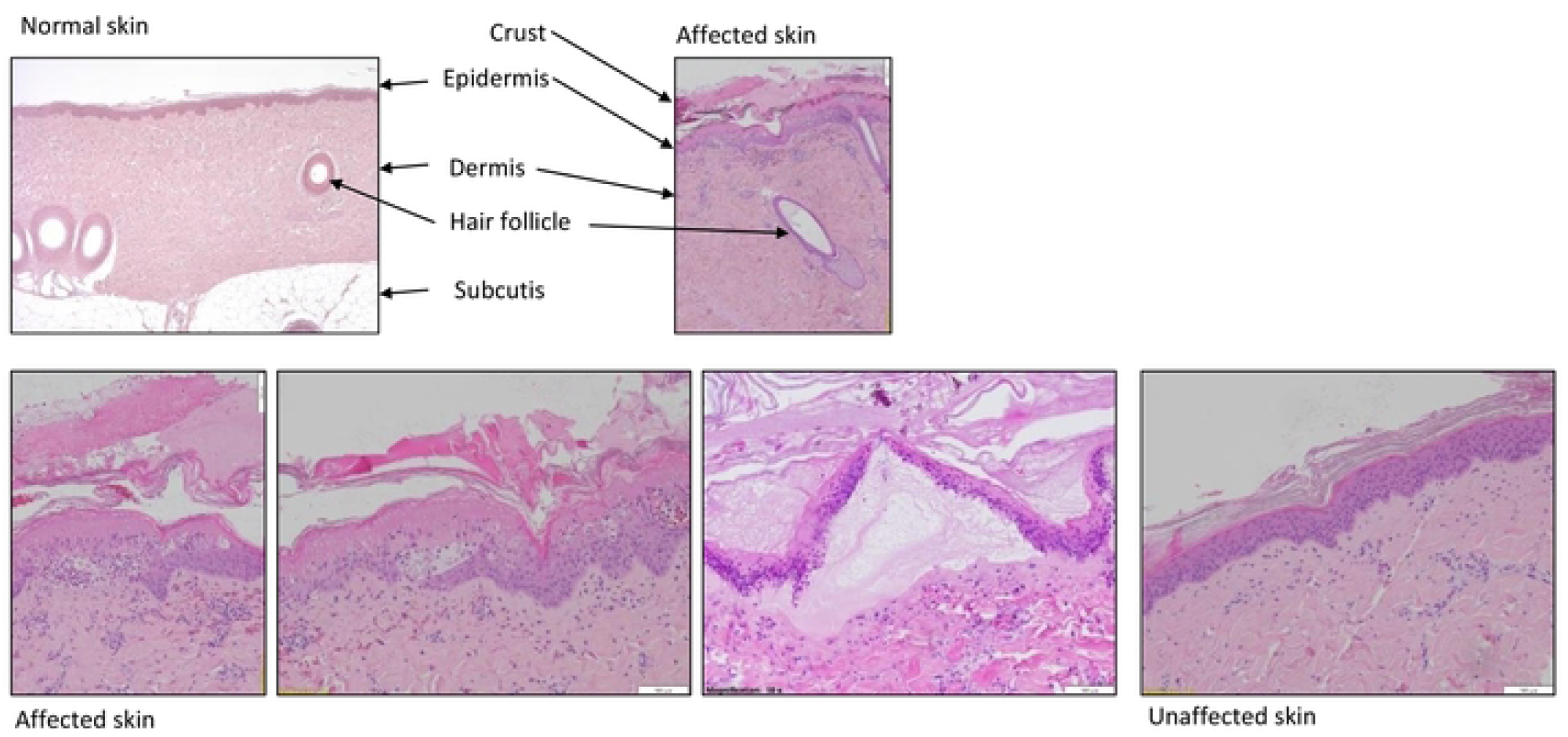
Histological analyses of normal, affected and neighboring unaffected skin. The pictures in the upper row had a magnification of 4x objective, that in the lower row of 10x objective.

In detail, in the affected skin of pig #239185, lesions were comprised by an acute inflammatory reaction, including epidermal necrosis, subepidermal edema, subepidermal infiltration of neutrophilic granulocytes and vesicle formation. Inflammatory lesions in affected skin of pig #314 also represented acute lesions, with epidermal necrosis, infiltration of mixed inflammatory cells (mainly granulocytes) in the epidermis or subepidermally, hemorrhage, intraepidermal pustule formation and subepidermal vesicles. Intercellular edema (spongiosis) was seen in the epidermis. Lesions in the affected skin of pig #901 were seen at the subepidermal/epidermal junction. Microscopic findings showed acute lesions, characterized by epidermal necrosis, epidermal intercellular edema (spongiosis), hemorrhage/congestion of blood vessels in the dermis, subepidermal vesicle formation and low-grade infiltrations of inflammatory cells (dominated by granulocytes). Lesions were seen at the subepidermal/epidermal junction. Lesions in the affected skin of pig #342036 were also acute and characterized by epidermal necrosis, epidermal intercellular edema (spongiosis) subepidermal infiltration of mixed inflammatory cells (dominated by granulocytes), hemorrhage/congestion of blood vessels in papillary dermis and subepidermal/dermal vesicle formation.

In one minipig (#237629), skin samples were obtained the day after clinical signs were observed. This minipig was treated against pain from the discovery of the clinical lesions until the animal was euthanized. Microscopic lesions were acute and similar to lesions in affected skin sampled from minipigs where skin was sampled on the same day when clinical lesions appeared. Lesions included epidermal necrosis, infiltration of mixed inflammatory cells with pustule formation and erosions/ulcerations. Pustule formation as well as erosions/ulcerations were all localized within or in relation to the epidermis or subepidermally.

Normal skin sampled from the affected animals did not show any pathological findings at the microscopic examination.

Skin was additionally sampled from one minipig (#342746) showing clinical signs of DPS 16 days prior to sampling. In the skin examined from this minipig, only parakeratotic hyperkeratosis was observed, consistent with healing of the lesions.

### No virus particles detected by electron microscopy

When an ultrathin section electron microscopy of skin samples was performed as described previously [45], no virus particles were detected, neither in the affected skin, nor in the unaffected skin.

### Selection of the porcine viruses to be tested for

Sensitive detection methods have been developed for use in the virological screening of genetically modified donor pigs generated for xenotransplantation as well as screening of the corresponding non-human primate transplant recipients [46]. These methods were also used to screen for viruses in GöMPs as these may serve as donor animals for islet cell transplantation into human diabetic recipients [23–29, 47]. Other pig breeds such as the Aachen minipigs [48, 49], the Mini-LEWE minipigs [32] and Greek pigs with erythema multiforme [16] were analyzed using these methods. The rationale for the selection of these so-called xenotransplantation-relevant porcine viruses [46] was among others the fact that HEV is indeed a well-known zoonotic virus [50] and PCMV/PRV has been shown to significantly reduce the survival time of pig xenotransplants in non-human primates [51–53]. PCMV/PRV was also transmitted in the first transplantation of a pig heart into a patient in Baltimore [54, 55] and it certainly contributed to the death of the patient since the clinical signs observed in the patient were the same as observed in baboons which received a PCMV/PRV-positive heart [51].

### Screening for porcine herpesviruses (PCMV/PRV and PLHV)

First, four porcine herpesviruses were included in the screening program, as erythema multiforme in humans is in many cases associated with a herpes simplex virus [11]. PCMV/PRV was detected in two minipigs, minipig #901 (liver, spleen and skin A, B, C) and in minipig #239185 (skin C). All other affected minipigs and the unaffected minipig were negative for PCMV/PRV (Figure 3, Supplementary Table 1). In minipig #237587 PCMV/PRV was not detected in PBMCs, even not after stimulation with the T-cell mitogen phytohemagglutinin (PHA), but it was detected in the liver, in four different liver lobes: left lateral lobe ct 38.28, left middle lobe ct 36.33, right middle lobe ct 35.58, right lateral lobe ct 37.42. The differences in the ct values indicate differences in virus replication in different liver regions.

**Fig 3.**
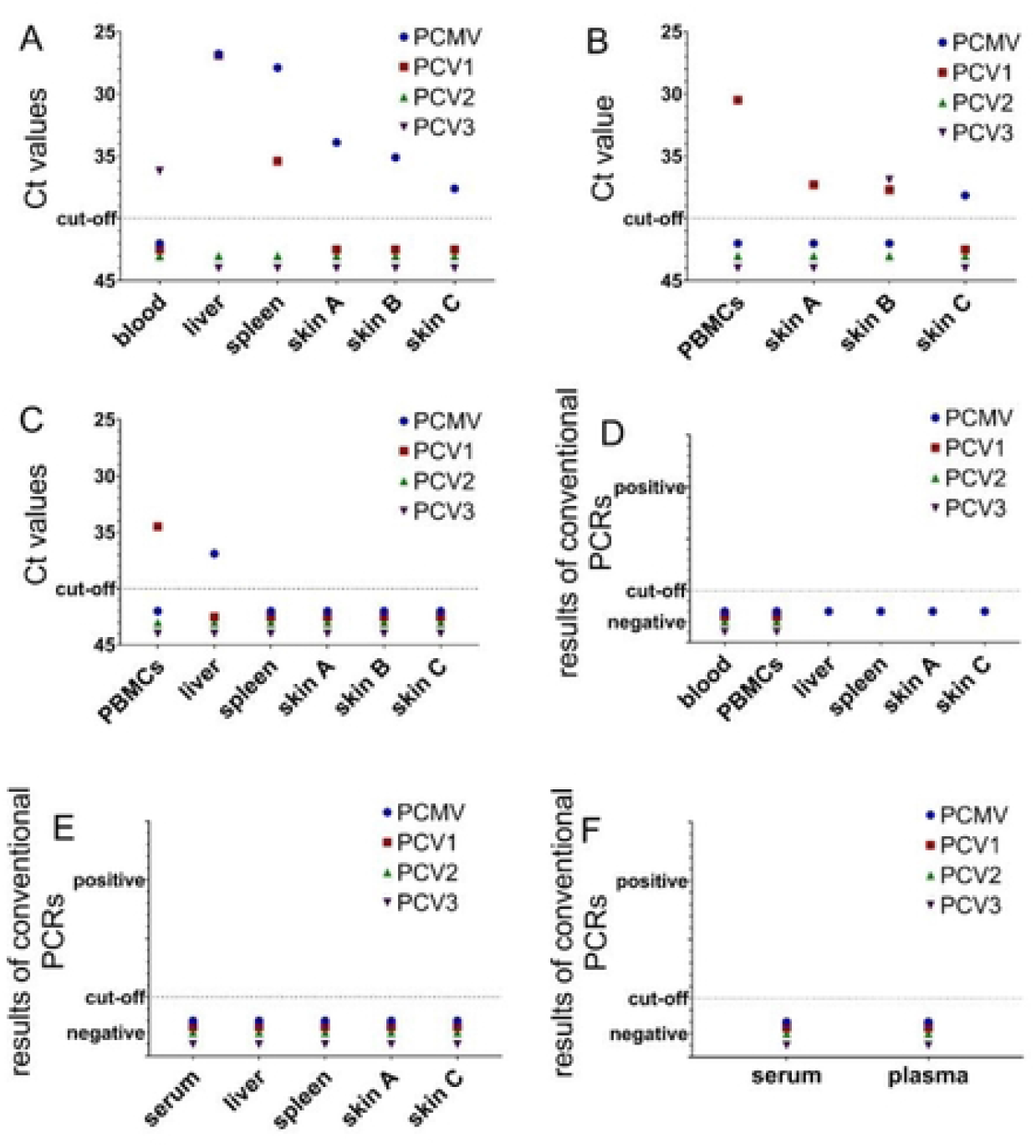
PCR results of the screening of six Göttingen Minipigs. Porcine viruses (PCMV, PCV1, PCV2, PCV3) were detected by real-time PCR and expressed as ct value (A-C) and conventional PCR expressed as detected/nondetected (D-F) in six minipigs suffering from dippity pig syndrome: pig #901 (A), pig #239185 (B), pig #237587 (C) and pig #342036 (D), pig #342746 (E), pig #343061 (F), whereas pig #343528 (C) did not show any clinical signs and were included as a reference? animal in this study.

Pig #349753 is the most thoroughly analyzed animal. From this animal sufficient material of the affected skin and different organs was available to perform all tests. In this animal, PLHV-3 was detected in the affected skin, in the border skin and in the unaffected skin and in other tissues (Table 3), suggesting that the virus was present also in all tissues of the animal. Also in this case, the differences in the ct values in different lobes of the liver indicate differences in virus replication in different liver regions. It was the only virus detected in this animal. It is important to note that PLHV-3 was not tested in some other animals with DPS due to the lack of sufficient material (Supplementary Table 1). All pigs were negative for PLHV-1, −2, and −4 (Supplementary Table 1). A summary of the results is given in Table 4.

**Table 3.** Screening for pig viruses in pig #349753

| Tissue | PCMV/<br>PRV | PLHV-1 | PLHV-2 | PLHV-3 | PPV-1 | PCV1 | PCV2 | PCV3 | PCV4 | PERVs | PERV-C | PERV-A/C | HEV |
| --- | --- | --- | --- | --- | --- | --- | --- | --- | --- | --- | --- | --- | --- |
|  | Real-time<br>PCR | Real-time<br>PCR | Real-time<br>PCR | Real-time<br>PCR | Real-time<br>PCR | Real-time<br>PCR | Real-time<br>PCR | Real-time<br>PCR | Real-time<br>PCR | Real-time<br>PCR | PCR | PCR | Real-time<br>RT-PCR |
| skin-affected area (SKA/A) | No ct | No ct | No ct | 35.31 | No ct | No ct | No ct | No ct | No ct | 12.23 | yes | no | No ct |
| skin non-affected area (SKNA/C) | No ct | No ct | No ct | 37 | No ct | No ct | No ct | No ct | No ct | 12.97 | yes | no | No ct |
| skin border between affected and non-affected area (SKB/B) | No ct | No ct | No ct | 38.43 | No ct | No ct | No ct | No ct | No ct | 13.15 | yes | no | No ct |
| Liver, left lateral lobe (LLL) | No ct | No ct | No ct | 37.23 | No ct | No ct | No ct | No ct | No ct | nt | yes | no | No ct |
| Liver, left medial lobe (LML) | No ct | No ct | No ct | 37.31 | No ct | No ct | No ct | No ct | No ct | nt | yes | no | No ct |
| Liver, right medial lobe (RML) | No ct | No ct | No ct | 39.65 | No ct | No ct | No ct | No ct | No ct | nt | yes | no | No ct |
| Liver, right lateral lobe (RLL) | No ct | No ct | No ct | 39.14 | No ct | No ct | No ct | No ct | No ct | nt | yes | no | No ct |
| Liver, caudate lobe (CAL) | No ct | No ct | No ct | 38.61 | No ct | No ct | No ct | No ct | No ct | nt | yes | no | No ct |
| Spleen (SPL) | No ct | No ct | No ct | 36.25 | No ct | No ct | No ct | No ct | No ct | nt | yes | no | No ct |

Nt, not tested, endogenous virus, present in all cells

**Table 4.** Summary of the virus detections in four Göttingen Minipigs with DPS and one animal not showing clinical signs using PCR methods. For details see Table 3 and Supplementary Table 3. The outstanding detection of PLHV-3 in animal 34753 is marked with a red circle.

| Animal | PCMV | PCV1 | PCV2 | PCV3 | PCV4 | PLHV-3 | PLHV-1,2,4 |
| --- | --- | --- | --- | --- | --- | --- | --- |
| 901 | + | + | - | + | - | nt | - |
| 239185 | + | + | - | + | - | nt | - |
| 237587 | + | + | - | - | - | nt | - |
| 343528 (No clinical signs) | - | + | - | + | - | nt | - |
| 349753 | - | - | - | - | - | + | - |

nt, not tested

Since PCMV/PRV is a herpesvirus capable of virus latency, analyses using PCR is not sufficient to detect the virus [44]. Therefore, serum and plasma from pig #349753 was tested in a Western blot assay for antibodies against PCMV/PRV, the result was negative (Figure 4).

**Fig 4.**
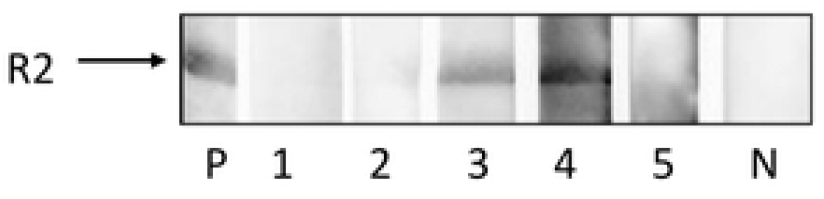
Western blot analysis of serum and plasma of pigs with and without DPS for antibodies against PCMV/PRV. Serum and plasma were tested at a dilution of 1:150 against the recombinant R2 fragment of the gB protein of PCMV/PRV, P, positive control serum; 1, serum #349753; 2, plasma #349753; 3, plasma #237587; 4, serum #901; 5, serum #343528 (without clinical signs), N, negative control serum. Exposure time 1 sec.

In the past, unaffected GöMPs from two facilities, from the Ellegard Göttingen Minipigs A/S and the Göttingen University, were screened for PCMV and PLHV-3 using real-time PCR [25–27] (Table 5). 12/39 animals (30%) were PCMV/PRV-positive in the first facility, none of ten animals in the second facility. PLHV-3 was found in two of 11 animals from the Göttingen University [47] (Table 5).

**Table 5.** Summary of the screening of control Göttingen Minipigs from Ellegaard Göttingen Minipigs A/S and from the University Göttingen using PCR-based and Western blot analysis. The references are indicated in brackets.

| Virus | GöMP from Ellegard Göttingen Minipigs A/S PCR method* | GöMP from the Göttingen University [44] PCR method* |
| --- | --- | --- |
| HEV | 9/40 [24] | 0/10 |
| PCMV/PRV | 12/39 [25, 41] | 0/10 |
| PLHV-1 | 1/20 [25, 41, 74] | 2/11 |
| PLHV-2 | 0/10 [74] | 2/11 |
| PLHV-3 | 0/20 [25, 74] | 2/11 |
| PCV1 | 0/21 [60] | n.t. |
| PCV2 | 0/21 [60] | 2/10 |
| PCV3 | 0/10 [unpublished] | 0/10 |
| PERV-C | 26/26 [44, 58] | 10/10 |

\*Positive animals/total animals using nested and/or real-time PCR; n.t., not tested

### Expression of PLHV-3

Since PLHV-3 was the only virus found in minipig #349753, its expression was analyzed using a real-time RT-PCR. Neither in the affected skin, nor in the unaffected skin, nor in the border skin, nor in liver lobes, nor in the spleen mRNA of PLHV-3 was found (not shown). This indicates that the virus is present in the animal, but it is replicating only below the detection limit of the used method.

### Screening for potentially zoonotic (HEV, SARS-CoV-2) and several other porcine viruses (PCV1,2,3, TTSuV and PPV1)

PCV1 was found in minipig #901 (liver and spleen) and minipig #239185 (PBMCs, skin A and skin B), but also in minipig #343528 not showing clinical signs (blood). PCV3 was found in minipig #901 (blood) and in minipig #239185 (skin B). PCV1 and PCV3, but no other viruses were found in GöMPs #901 and #343528, which were housed closely together. In conclusion, minipig #901 was infected with three viruses (PCMV/PRV, PCV1 and PCV3) and minipigs #239185 and #237587 with two viruses (PCMV/PRV and PCV1 or PCV1 and PCV3). In all other animals none of these three viruses were detected. The viruses PCV2, PCV4, PPV1, TTSuV1, TTSuV2 and HEV were not found in any of the tested animals. In previous studies analyzing GöMP PCV2 was found in five of 31 animals [27, 29] and HEV in nine of 40 animals [24] (Table 5). SARS-CoV-2 was not found in the pigs. This is in concordance with extensive studies showing that pigs cannot be infected with this virus [56–59]. All these viruses were also not found in pig #349753 (Table 3).

### Prevalence of PERV-C

It is well known, that PERV-A and PERV-B are present in the genomes of all pigs, whereas PERV-C is not necessarily present in all pigs. Therefore, all animals with DPS and the animal with no clinical signs were screened for PERV-C. All animals had integrated PERV-C (as well as PERV-A and PERV-B) in their genome (Table 3, Supplementary Table 4). This is in concordance with previous investigations showing that all GöMPs carry PERV-C in their genome [27, 28] (Table 5).

### Enhanced expression of PERV

When the expression of PERV was investigated using a real-time RT-PCR, expression in minipig #901 was detected in the liver and spleen, no expression was detected in the affected and unaffected skin (Figure 5A). In minipig #237587 expression was detected mainly in unstimulated and stimulated PBMCs and to a lower extend in liver lobes and spleen detected in the liver (Figure 5C). It is important to underline that PERV expression is enhanced in PBMCs from pigs #239185 stimulated by the T cell mitogen phytohemagglutinin (PHA) compared with unstimulated PBMCs (Figure 5D), confirming previous results in GöMPs [27, 28, 44] and other pig breeds [60] that treatment with a mitogen, here PHA, increases PERV expression.

**Fig 5.**
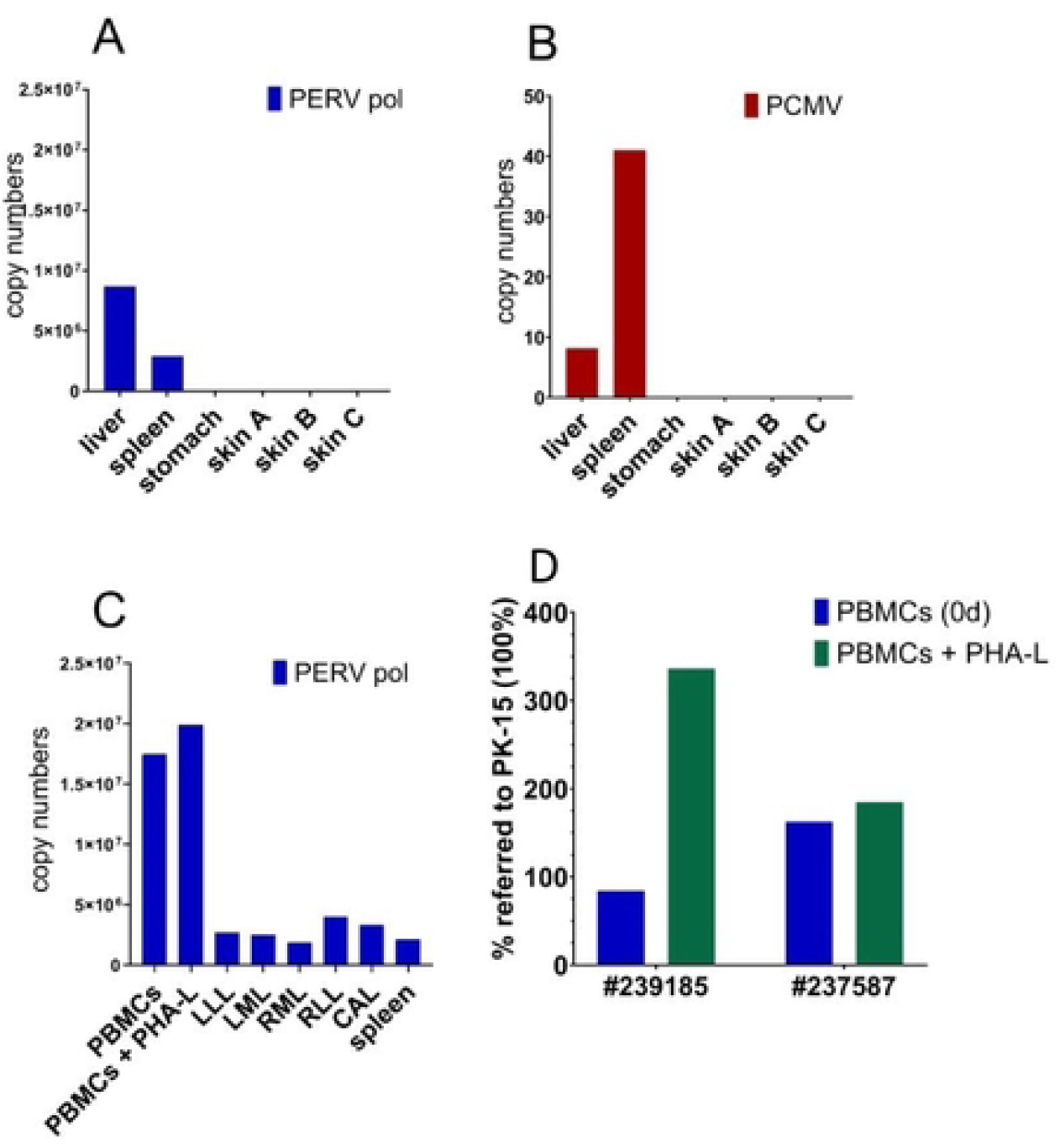
Gene expression of PERV and PCMV in different organs of pigs with DPS. Gene expression is displayed for two minipigs: minipig #901 (A and B) and minipig #237587 (C). Skin A, B, C refer to the skin regions indicated in Figure 1. PHA-L, lymphocyte-specific phytohemagglutinin. The liver lobes are indicated as: LLL, left lateral lobe; LML, left medium lobe; RML, right medium lobe; RLL, right lateral lobe; CAL, caudate lobe. PERVpol indicated primer pairs used to detect the highly conserved polymerase gene of PERV. D, Increase of PERV expression after mitogen stimulation. PBMCs from pig #239185 and pig #237587 were incubated with the mitogen lymphocyte-specific phytohemagglutinin (PHA-L). Gene expression of PERVpol was referred to the expression in PK-15 cells which was set 100%.

### Detection of PERV-A/C

PERV-A/C recombinants are the product of a recombination between PERV-A and PERV-C, the recombinants can be found in somatic cells, but never in the germ line of pigs. Since in all analyzed minipigs PERV-C was found integrated in their genome, the probability of recombination with PERV-A exists. Among the minipigs with DPS, PERV-A/C was found only in one animal, #342036 (Supplementary Table 1). PERV-A/C was found in the liver, spleen, in the affected and also in the unaffected skin of this animal. PERV-A/C was also found in freshly isolated PBMCs and in PBMCs 5 days after incubation with and without PHA-L. In the cultured PBMCs the expression of PERV-A/C was 13 times higher compared with the uncultured, in the PHA-stimulated PBMCs 42 times higher compared with the expression in unstimulated PBMCs. Previously, PERV-A/C was not detected in the germline of 26 GöMPs from Ellegaard Göttingen Minipigs A/S in two studies [27, 28], however it was found in the genome of freshly isolated PBMCs from two GöMPs from the Göttingen University [27] and was released from PBMCs of one animal and infected human cells [27, 28].

### Expression of PCMV/PRV

PCMV/PRV was detected in the liver, spleen and all skin samples (skins A, B and C) of minipig #901. Whereas PCMV/PRV specific mRNA was found in liver and spleen, the expression of PCMV/PRV was lower in the skin samples (Figure 5B). When serum from animal #901 was screened for antibodies against PCMV/PRV performing a Western blot analysis using a recombinant C-terminal fragment of the gB protein of PVMV/PRV, virus-specific antibodies were detected, indicating the expression of virus protein and antibody production (Figure 4). In contrast, sera or plasma from animals #349753 and #343528 were negative in the Western blot analysis (Figure 4). Animals #349753 and #343528 (the animal without clinical signs) were negative in the PCR assay (Table 3, Table 4) Serum from animal #237587 was positive for antibodies against PCMV/PRV and in the PCR assay (Supplementary table 1, Figure 4).

### Results of the next generation sequencing

A differential gene expression was observed when skin punches from affected (skin A) and unaffected skin (skin C) from pig #237587 were analyzed for gene expression on an Illumina platform, (Table 6, Supplementary Table 2, Supplementary Table 3). Significant differences were observed when the bacteria where analyzed (Table 6). Concerning the viruses, an elevated expression of retroviruses was observed (0.20169% in the affected skin versus 0.14270% in the unaffected skin). Viruses of the families *Bromoviridae* (single stranded DNA viruses), *Partitiviridae* (double stranded RNA viruses), and *Totiviridae* (also double stranded RNA viruses), which are all three increased in the affected skin, are viruses infecting plants, fungi and parasites. They obviously are associated with the hay or straw in the animal stables.

**Table 6.**
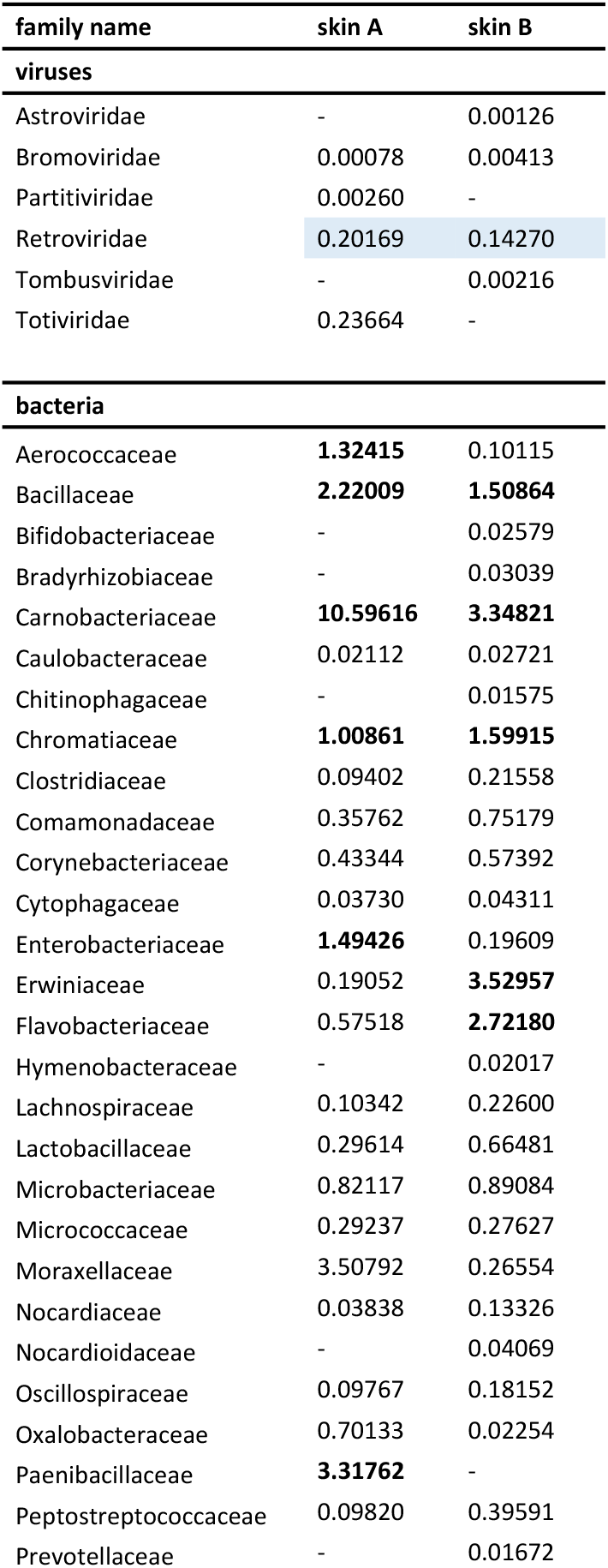
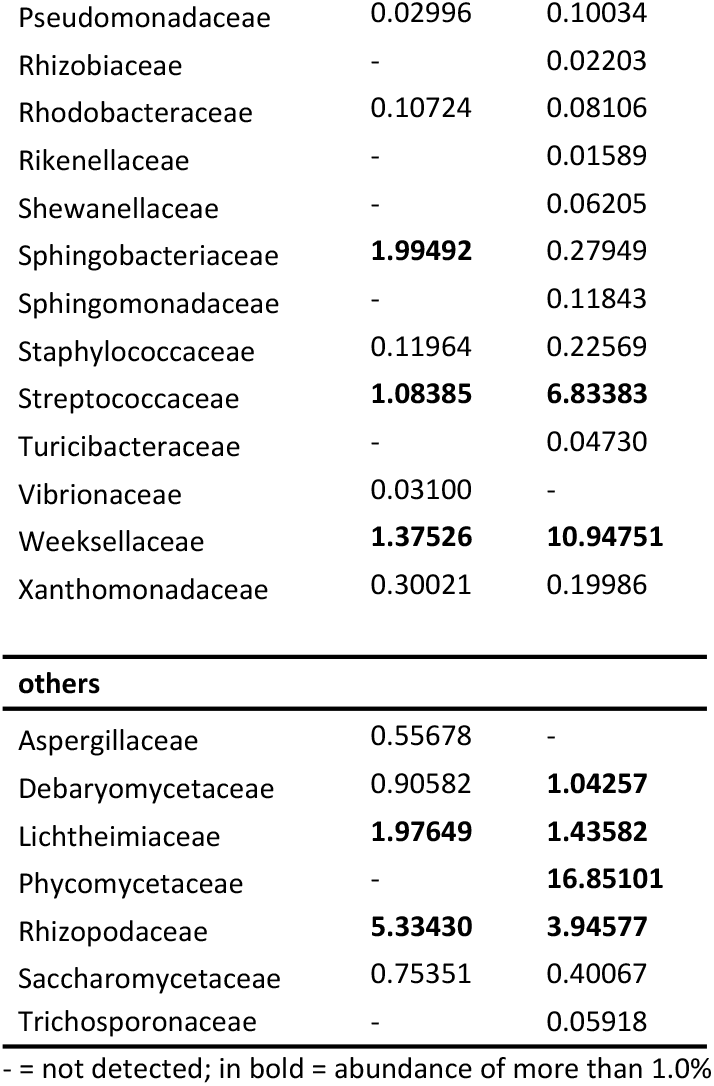
Comparative analysis of the abundance (in %) of taxonomic families obtained by a NGS approach for two skin areas from animal 237587. Skin A = affected region, skin B = not affected region.

## Discussion

This is the first comprehensive analysis of GöMP with DPS, searching for a virus which may be the cause or at least involved in the pathogenesis of DPS. Since the DPS has similarities with erythema multiforme in pigs and humans and since the human erythema multiforme is associated with a herpesvirus, HSV, a herpes virus was suggested to be involved in the pathogenesis of DPS. In addition, the pig syndrome is only of short duration similarly as herpes labialis lip injury in humans, Most importantly, in the best analyzed GöMP, minipig #349753, only one virus was detected in the animal, PLHV-3 (Table 3). PLHV-3 could not be analyzed in the pigs #901, and #239185, due to restricted sample material for testing. PCMV/PRV was found in GöMPs #901, #239185, and #237587, (Supplementary Table 1), but not in minipig #349753, neither by PCR nor by antibody testing.

Most interestingly, GöMPs #901, #239185, and #237587 with DPS were positive for PCV1, but the GöMPs without clinical signs, #343528, was PCV1-positive, too. Minipig #343528 was housed at Scantox and not in the breeding colony of Ellegaard Göttingen Minipigs A/S. The detection of PCV1 in GöMPs is something new. When GöMPs from the Ellegaard Göttingen Minipigs A/S facility were screened the first time, the animals were not tested for PCV1 [24, 25]. When they were tested for PCV1 the first time, none of the 21 screened animals were positive [29]. The minipigs with DPS #342036, #342746, #343061, tested still at RKI, were also PCV1-negative (Supplementary Table 1). However, the testing was based on a conventional PCR which is less sensitive compared with the more sensitive real-time PCR. The real-time PCR was used for testing the minipigs #901, #239185, #237587 and #343528, at the Institute of Virology. PCV1 was thought to be apathogenic in pigs (for review see [61, 62]). Since the minipig not showing any clinical signs was also PCV1-positive, this virus could contribute to the DPS only when another factor is involved. Such a co-factor may be stress, bacterial infection or infection with another virus.

Screening for viral infections was complicated by difficulties in isolating DNA and RNA from pig skin, especially from the affected skin. The skin structure of mammals is mainly composed of three distinct layers: subcutis, dermis, and epidermis. Pig skin has a similar epidermis as human skin, with a comparable thickness. Both human and porcine dermis is divided into a papillary layer (pars papillaris) and a reticular layer (pars reticularis). Porcine dermal collagen is similar to human dermal collagen biochemically. The subcutis is the fatty subdermal layer of the skin. In humans and pigs, this fat layer is the main insulation component [63]. Therefore, several histological and immuno-histochemical similarities exist between porcine and human skin [64]. With a few exceptions, there are no publications describing how to isolate nucleic acids from pig skin [30, 65]. Based on these publications we included an incubation step of the skin with collagenase and hyaluronidase (see Materials and methods).

It is of great interest, that PERV-A/C was found in one of the DPS pigs, pig #342036. PERV-A/C recombinants are characterized by higher replication titers compared with the paternal PERV-A [66, 67]. The virus was found in different organs of the pig, including the affected and unaffected skin. PERV-A/C was highly expressed in the PBMCs after incubation with and without PHA-L. This is also new since in previous investigations PERV-A/C was never found in GöMPs from the Ellegaard Göttingen Minipigs A/S facility [28]. PERV-A/C was however found in two cases of GöMPs from the facility of the Göttingen University, were the animals were developed in the past [27, 47]. In one case the newly isolated recombinant PERV-A/C was able to infect human cells [27, 47]. Whether the recombination and the wide distribution of PERV-A/C in minipig #342036 is associated with the DPS features remains unclear.

Performing next generation sequencing (NGS) we expected additional information on putative virus sequences in the affected skin region in comparison with the unaffected skin. However, the results showed neither new porcine viral sequences, nor a difference between affected and unaffected skin regions concerning pig viruses. As known from different analyses of the pig virome, viruses with known or suspected zoonotic potential were often not detected by NGS, but were revealed by more sensitive PCR-based methods (for review see [68]). Our data confirm these findings: Viruses detected in the skin of minipig #237587 by PCR, PCMV/PRV and PCV1 (Figure 2, Table 4) were not detected by NGS, with exception of the retroviruses, which were expressed higher in the affected skin compared to the unaffected skin (Table 3). Therefore, this method did not contribute to the solution of the problem, which virus, if any, may be involved in the pathogenesis of DPS in this case.

Only PLHV-3, but no other viruses, was found in one animal, minipig #349753. It was found in the affected skin, but also in the unaffected skin and other organs of the pig (Table 3). It is important to note that PLHV-3 was also found in four of five Greek pigs suffering from erythema multiforme [16]. Unfortunately, animals #901, #239185, and #343528 were not tested for PLHV-3 due to the absence of sufficient material. PLHV-3 was not found in previous studies in GöMP from the Ellegaard Göttingen Minipigs A/S facility [25, 69], but 2 out of 11 animals from the Göttingen university facility were positive [27] (Table 6).

PLHV-3, also known as suid herpesvirus 5 (SuHV-5), is a gammaherpesvirus assigned to the genus Macavirus [70]. The role of PLHV-3 as primary pathogen of swine, or as co-factors in other viral infections, is largely unknown. PLHV-3 was frequently found in the blood and in lymphoid organs of domestic and feral pigs from different geographic locations. PLHV-3 was detected predominantly in B-cells [34]. 48% of German pigs and 65% of Italian pigs were found PLHV-3 positive [34]. The prevalence of PLHV-3 varies between 5 and 65% in various studies performed in Germany, Italy, Spain, France, USA, and Ireland [34, 39, 72, 73]. PLHV-3 was also detected in wild boars [74]. A latent as well as productive PLHV-3 infection was found in the porcine B-cell line L23 [34, 71]. Since PLHV-3 is a herpesvirus capable of latency, a Western blot assay was developed based on a recombinant fragment of the gB protein of PLHV-1, which also should detect antibodies against PLHV-3 [69]. Using this newly developed Western blot analysis, PLHV was detected in slaughterhouse pigs [69], and in Aachen minipigs [75], but not in GöMPs [69]. Using a PCR method, PLHV-3 was found in 0/10 GöMPs, 2/8 Aachen minipigs, 0/10 Mini LEWE minipigs [32] and 10/36 slaughterhouse pigs [69]. The fact that PLHV-3 was not found previously in GöMPs using PCR [25, 27, 69] is interesting considering that PLHV-3 was detected in the affected skin, the unaffected skin and other organs of the GöMP #349753 suffering from DPS (Table 3). The fact that all viruses found in DPS pigs were also found in animals without symptoms of disease, suggests a multifactorial cause of DPS.

Of interest is also the fact that the expression of PERVs is slightly higher in the affected skin compared with the unaffected skin (Table 6). This may be due to inflammation and infiltration of activated immune cells. The expression of PERVs is much higher in PBMCs after stimulation by the mitogen PHA, simulating the activation of immune cells in an immune response as shown by RT-PCR (Figure 5D)

Further investigations on minipigs suffering from DPS should be performed. First of all, more tissue samples are needed, especially from the affected and neighboring skin. However, samples from other tissues are also required for comparative analyses. This will allow screening for all viruses and performing repeated testing.

To summarize, we detected different viruses in the affected animals and in the affected skin, with one individual being of great interest due to having only PLHV-3. However, all viruses have also been found in unaffected animals. Consequently, we suggest that DPS has a multifactorial cause. However, elimination of the detected viruses from GöMPs using early weaning, vaccines, antiviral drugs, Caesarean delivery, colostrum deprivation, and embryo transfer may prevent DPS.

## Supporting information

**Supplementary Table 1.** Overview of the PCR results of six Göttinger Minipigs with dippity pig syndrome (DPS).

**Supplementary Table 2.** Abundance of microbiological taxa obtained by an NGS approach for skin A (affected skin region) from animal 237587. The selected taxa refer to the least common denominator of the data revealed by NGS approach independently of the taxon.

**Supplementary Table 3.** Abundance of microbiological taxa obtained by an NGS approach for skin B (affected skin region) from minipig #237587. The selected taxa refer to the least common denominator of the data revealed by NGS approach independently of the taxon.

**Supplementary Table 4** Detection of integrated PERVs in six Göttingen Minipigs with DPS and one unaffected animal.

## Acknowledgement

We wish to thank the caretakers, technicians, pathologists and veterinarians involved in the sampling and handling of tissue from the affected animals. The research in the laboratory of Joachim Denner at the Robert Koch Institute and the Institute of Virology was supported by the Deutsche Forschungsgemeinschaft, TRR127.

## Authors contribution

**Conceptualization:** Joachim Denner, Kari Kaaber

**Resources and analysis:** Kari Kaaber; Nanna Grand, Kirsten Rosenmay Jacobsen, Michelle Salerno

**Virological analysis:** Hina Jhelum, Sabrina Halecker, Robert Prate, Luise Krüger, Yannik Kristiansen, Ludwig Krabben, Benedikt Kaufer, Joachim Denner

**Electron microscopy:** Lars Möller, Michael Laue

**Histological analysis:** Nanna Grand

**Writing – original draft:** Joachim Denner, Kari Kaaber

**Writing – review and editing:** Hina Jhelum, Nanna Grand, Kirsten Rosenmay Jacobsen, Michelle Salerno, Sabrina Halecker, Ludwig Krabben, Benedikt Kaufer, Kari Kaaber, Joachim Denner

